# Low-cost platform for multi-animal chronic local field potential video monitoring with graphical user interface (GUI) for seizure detection and behavioral scoring

**DOI:** 10.1101/2022.07.14.500102

**Authors:** Gergely Tarcsay, Brittney Lee Boublil, Laura A. Ewell

**Affiliations:** Anatomy & Neurobiology, 835 Health Sciences Road, University of California, Irvine 92697

## Abstract

Experiments employing chronic monitoring of neurophysiological signals and video are commonly used in studies of epilepsy to characterize behavioral correlates of seizures. Our objective was to design a low-cost platform that enables chronic monitoring of several animals simultaneously, synchronizes bilateral local field potential and video streams in real-time, and parses recorded data into manageable file sizes. We present a hardware solution leveraging Intan and Open Ephys acquisition systems and a software solution implemented in Bonsai. The platform was tested in 48-hour continuous recordings simultaneously from multiple mice (male and female) with chronic epilepsy. To enable seizure detection and scoring, we developed a graphical user interface (GUI) that reads the data produced by our workflow and allows a user with no coding expertise to analyze events. Our Bonsai workflow was designed to maximize flexibility for a wide variety of experimental applications, and our use of the Open Ephys acquisition board would allow for scaling recordings up to 128 channels per animal.

**SIGNIFICANCE STATEMENT:** We present a low-cost hardware and software solution intended for multi-animal chronic seizure monitoring, that prioritizes experimental freedom, and requires no coding expertise of the user. We provide details for using an Intan adapter board to enable user freedom regarding the type of electrodes used. Video and local field potential data streams are synchronized and parsed in Bonsai – an open-source visual programming language that has pre-written libraries that allow our workflow to be adapted to other data types or to integrate with open-source toolboxes. Finally, for those intending to use our platform for seizure monitoring, we provide an accessible GUI to aid in seizure detection and behavioral scoring.

## INTRODUCTION

Rodent studies of epilepsy often utilize longitudinal recordings to characterize disease progression and determine the efficacy of drug treatments (Price et al., 2009; reviewed in Simonato et al., 2014; Li et al., 2021; Lisgaras & Scharfman, 2022). In such experiments it is beneficial to simultaneously record video and electrophysiological signals such as electroencephalogram (EEG) or depth local field potential (LFP). Video-neurophysiology recordings allow for assessing the behavioral stage of seizure events recorded. Common practice is to score seizures using the Racine scale – a scoring system based on distinct behavioral phenotypes such as head movements, posture, forearm clonus, rearing, and falling (Racine, 1972). Categorization of the behavioral severity of seizures is critical for determining disease stage – for example, chronic epilepsy is defined by the occurrence of two or more behavioral seizures. In addition to studies of epilepsy, video-neurophysiology recordings are becoming widely used in rodent studies of other pathologies that are sometimes comorbid with epileptiform activity including depression (Sadeghi et al., 2021), Alzheimer’s Disease (Harris et al., 2010; reviewed in Vossel et al., 2017), and autism (Lewis et al., 2018; Eisenberg et al., 2021; reviewed in Specchio et al., 2022).

Off the shelf acquisition systems for long term video-neurophysiology recordings are costly and do not provide the user with state-of-the art hardware for *in vivo* neurophysiology. Despite this, such systems are appealing to new users because they are plug and play, automatically align video and neurophysiological data, and typically are packaged with analysis software. However, they often have relatively low channel counts with low sampling rates – and thus are only suitable for classic video EEG / LFP. For example, they would not be conducive for high density linear probe recordings or tetrode recordings which facilitate population and single unit recordings. Over the last decades, low cost, high channel count, high sampling rate data acquisition boards and miniaturized amplifiers have become increasingly available through companies like Intan and Open Ephys (Siegle et al., 2017). Large channel count systems (>256 channels) could theoretically support both low density electrophysiology in several animals (as is common in video-neurophysiology recording systems) or high-density electrophysiology in individual animals. With such systems, labs could invest in one acquisition system that would facilitate several types of experiments. These systems offer the low-cost flexible solution needed but tailoring them to maximize their flexibility requires significant knowhow. For high throughput video-neurophysiology monitoring several challenges need to be overcome by the user, such as (1) how to integrate multiple headstages to facilitate high throughput multi-animal recordings, (2) data synchronization between video and voltage data streams, (3) parsing large data files, and (4) analysis pipelines for systematically organizing and evaluating large datasets.

Here we provide a novel method for converting a single Open Ephys acquisition board to a high throughput video-local field potential (video-LFP) acquisition system. We establish a method for continuously recording from four mice simultaneously. We developed a software solution for synchronizing video and LFP streams, and for saving data files periodically to output data packets in sizes that are manageable for further processing. Lastly, we offer a home written graphical user interface (GUI) for automatically detecting putative seizures and facilitating easy annotation of correlative video behavior by the user.

## METHODS

### Subjects

14 C57BL/6 mice (six males and eight females), all aged between 10 to 12 weeks, were used for this study. Mice were housed individually on a reversed 12 h light/dark cycle and weight maintained on an ad libitum diet for the duration of the experiment. All procedures were approved by the University of California, Irvine Institutional Animal Care and Use Committee (IACUC).

### Custom-made electrodes

Electrodes were constructed from perfluoroalkoxy alkane (PFA) coated, stainless steel wire (127 μm bare, 203.2 μm coated, SS-5T, Science-Products, Germany) and gold plated Mill-Max male pins. Insulation was removed from one end of the wire and then wrapped around and soldered to the base of the wrap-post of the pin. The wrap-post was trimmed with wire cutters and the junction was insulated with epoxy. Finally, the wire was trimmed to the desired length (∼ 2 mm) based on the coordinates for the target brain region (Figure 1 A). Impedance measurements ranged from 100 – 200 kOhm. For ground electrodes the procedure was the same except the insulation was removed from the entire portion of the electrode that would be inserted into the brain, resulting in significantly lower impedance.

**Figure 1.**
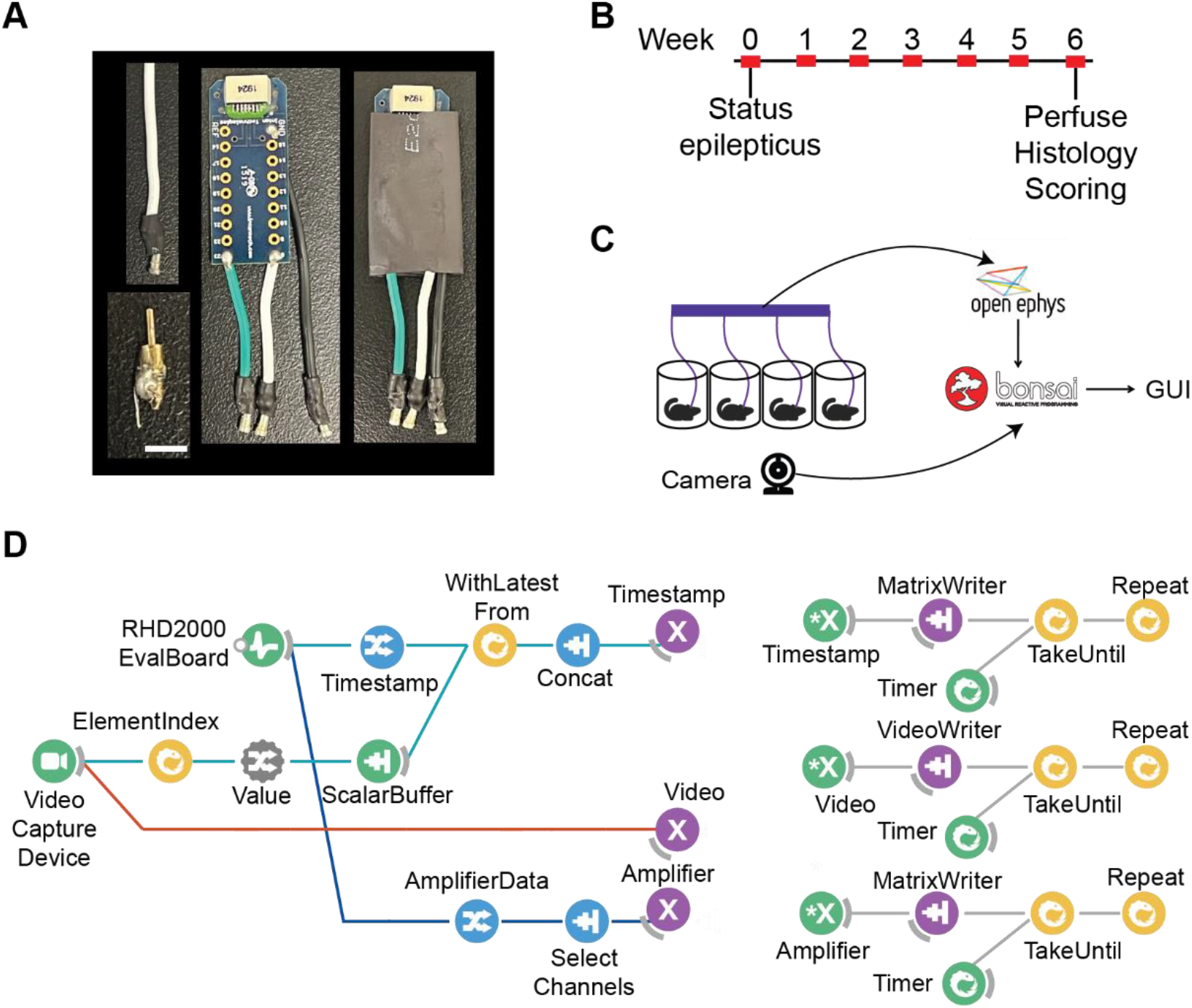
Schematic of multi-animal chronic video-LFP recording and analysis pipeline. **A** Components for chronic LFP recordings. Custom-made electrodes (bottom left, scalebar: 3 mm) were implanted into CA1 of the hippocampus. Custom-made cables (top left) were soldered to an Intan adapter board (middle) and further protected with 10 mm heat shrink tubing (right). **B** Timeline of experiment. Red rectangles represent 48 hours of continuous monitoring. **C** Experimental setup. Recordings were performed from 4 mice simultaneously. Video-LFP data was acquired and controlled in a custom-made Bonsai workflow. Seizure analysis was performed with the custom-made Seizure Analysis GUI, written in MATLAB. **D** LFP and video were acquired and synchronized by merging timestamps (left). Timestamp, video, and voltage data were streamed to time-dependent loops (nodes labelled with X and *X) that enabled data saving once per hour.

### Wiring the Intan adapter board

Custom cables were made from 26-gauge PVC wire (BNTECHGO, USA) and secured to an Intan 18-pin electrode adapter board (Part Number C3418). For ease of identification during experiments, we recommend using a different color for each cable. For our experiment, we used black for the ground, white for the right electrode, and green for the left electrode. To assemble the cables, a cm of insulation was removed from both ends of each wire. Next, one end of exposed wire was soldered to the base of the wrap-post of a female Mill-Max pin. A small piece of 2.5 mm diameter heat shrink tubing (Amazon, B08N5246YB) was placed around the solder joint between the wire and pin for additional support and insulation. The remaining end of the wire was soldered to one of the channels on the Intan adapter board. Once all cables were constructed and attached to the adapter board, we wrapped the board in 10 mm heat shrink tubing to give additional support and protection of the cables (Figure 1 A).

### Surgical procedures

All surgeries were performed using aseptic procedures. Anesthesia was maintained throughout the surgery with isoflurane gas (1%-2% isoflurane delivered in O_2_ at 1 L/min), and carprofen (5 mg/kg, Zoetis Inc., USA) was administered subcutaneously as an analgesic. Animals were positioned in a Kopf stereotaxic instrument with non-rupture ear bars (David Kopf Instruments, USA). Body temperature was maintained at 37-38 °C using air-activated heating packets (HotHands, USA). Hair was removed from the scalp and the skin was washed with iodine (Dynarex Corporation, USA). The skin above the skull was injected with lidocaine (4 mg/kg, Sparhawk Laboratories Inc., USA). A midline incision was made in the scalp and the skin was retracted to expose the skull. The skull bone was prepared and cleaned using a scalpel and sterile cotton swabs.

Using a spherical carbide bur (HP ¼, SS White Dental, USA), shallow divots were made over the skull’s surface to create additional surface area to improve bonding with light activated glue. A thin layer of Optibond (Kerr Italia, Italy) was applied over the skull and cured for 30 seconds with UV light (Superlite, MW Dental, Germany). Three craniotomies were made using the same spherical carbide bur, one over the left and right hippocampus (AP: −2.0 mm, ML: ±1.5 mm from bregma) and one over the cerebellum behind lambda for the ground electrode. If bleeding occurred, it was immediately stopped with Gelfoam (Ethicon Inc., USA). Saline (Teleflex Medical, USA) and Gelfoam were placed over each craniotomy to ensure the brain tissue remained protected and healthy. Custom-made electrodes (described above) were first implanted into the hippocampus contralateral to the side where we would eventually inject kainic acid (see below) (DV: −1.3 mm from dura) and into cerebellum to serve as the ground. All electrodes were secured to the skull using an optical dental composite (Tetric EvoFlow Ivoclar Vivadent, Switzerland).

To induce *status epilepticus*, we adopted the supra-hippocampal kainic acid model of chronic epilepsy (Bedner et al., 2015). Kainic acid (KA) (Tocris, USA) was dissolved in saline to provide a solution with a concentration of 20 mM, pH 7.4 (Scharfman, Goodman, & Sollas, 2000; Iyengar et al., 2015; Lisgaras & Scharfman, 2022). KA was injected supra-hippocampally using a custom microinjection system (Zhu, Eom, & Hunt, 2019) (MO-10, Narishige International, USA). The glass pipette was lowered into the brain (DV: −0.8 mm from dura) and left in place for a minimum of 1 minute before beginning to inject. 70 nL of KA was injected over the span of two minutes. The glass pipette remained in place for 1 minute following the injection to reduce the spread of the drug up the needle tract. Following the injection, the remaining electrode was implanted and secured using the same procedures described above. At the completion of the surgery, saline was administered subcutaneously. *Status epilepticus* was terminated 4 hours after the completion of the surgery with a subcutaneous injection of lorazepam (7.5 mg/kg, West-Ward, USA). All animals received postoperative care for at least 7 days after surgery and daily monitoring of weight and overall health continued for the duration of the experiment.

### Data acquisition

The adapter board was connected to an Intan RHD 16-channel headstage (Part Number C3334) that sends the digitized signal to the Open Ephys acquisition board (Siegle et al., 2017) via an Intan RHD 6-ft ultra-thin SPI interface cable (Part Number C3216). Connections at the subject’s head were wrapped with beeswax (Amazon, B07VSQ6PGD) to prevent detachment. Local field potentials (LFP) were sampled at 2000 Hz and filtered in the 0.1-800 Hz range. Data was transmitted to a HP Z2 G5 TWR PC through a USB 3.0 connection. We were able to monitor four mice simultaneously as the acquisition board is equipped with four headstage SPI connectors. Electrical and mechanical noise were minimized by a custom-built Faraday-cage and grounding of both the Faraday-cage and the ground screw terminal of the acquisition board (accessible on the side of the Open Ephys box). Mice were monitored and video recordings were obtained using a 30 Hz Spedal 920 Pro wide angle web camera (Amazon, B07TDQ8NL3).

### Monitoring sessions

LFP and video were monitored for 48 hours per week for 6 weeks per animal (Figure 1 B). After surgery, animals were immediately connected to the acquisition system in order record *status epilepticus*. Cages were illuminated with red and white light to maintain the reversed 12 h light/dark cycle. Red light is not in the visual spectrum of mice, so allowed for ‘darkness’ while providing light for video monitoring. Mice were provided with water-soaked food and a non-wetting water gel (HydroGel, ClearH_2_O, USA) daily during the monitoring sessions. At the end of each 48-hour monitoring session, pins were unplugged, and mice were transferred back to their home cages.

### Bonsai workflow and data synchronization

A data workflow was developed in Bonsai, which is an open-source visual programming software for collecting, processing, and controlling distinct data streams (Lopes et al., 2015; https://bonsai-rx.org/). Data are represented as observable sequences, such as series of measured voltage values or acquired camera frames. In Bonsai, observable sequences are acquired and controlled by reactive operators, that are visualized as colored nodes. Each operator’s behavior can be configured by setting its parameters in the Properties panel. *Source operators* are used to generate observable sequences (such as acquiring data from a device or generating an integer). *Transform operators* modify the observable sequences (such as filtering). *Combinators* can be used to merge distinct data streams. *Subject operators* are used for reusing and sharing observable sequences. Processed data streams are saved by *sink operators*.

Necessary operators for LFP acquisition and control are provided in the Bonsai Ephys library, while the camera is accessed through an operator in the Bonsai Video library. LFP and video were visualized in real-time during each monitoring session with Bonsai Dsp Design Library. LFP and video data were saved by operators of the Dsp and Vision libraries, respectively. Asynchronous video and LFP sequences were synchronized by chaining together *combinator operators* that merge the latest video timestamp and the timestamp of the currently acquired LFP sample resulting in a two-column vector. Our workflow can be found at https://github.com/EwellNeuroLab/Chronic-Recordings-Bonsai-Workflow. Visual examples of all property settings can be found at https://github.com/EwellNeuroLab/Chronic-Recordings-Bonsai-Workflow/tree/main/Bonsai%20Property%20Settings.

#### LFP acquisition in Bonsai

LFP data from the Open Ephys board was acquired by the *RHD2000EvalBoard* source operator. The *RHD2000EvalBoard* source has several parameters which can be set by the user including sampling rate (2000 Hz), analog high and low-cut filters on the Intan chip (0.1 – 800 Hz), and a DSP high cut filter (1 Hz). An Open Ephys bit file must be provided for FPGA configuration to allow communication between Bonsai and the acquisition hardware (https://github.com/open-ephys/GUI/tree/master/Resources/Bitfiles) and should be renamed as ‘main.bit’ to be compatible with our workflow. Furthermore, the *RHD2000EvalBoard* source has several data fields, such as ‘Timestamp’ and ‘AmplifierData’. In order to extract them individually, data fields can be selected in the Output menu of the *RHD2000EvalBoard* source. After selection, a *MemberSelector* operator is created that maps the individual data field from the source node. In order to extract the voltage data, *AmplifierData* MemberSelector was created by selecting the ‘AmplifierData’ data field and passed to the *SelectChannels* filter that allowed us to select the channels of interest.

#### Synchronizing LFP and video data streams

We created the *Timestamp MemberSelector* operator by selecting the ‘Timestamp’ data field of the *RHD2000EvalBoard* source and timestamps of the video frames were accessed via the *ElementIndex* combinator (Figure 1 D, left). The Open Ephys acquisition board buffers 256 sampling points of voltage data and sends timestamp in 32-bit integer format. Therefore, the timestamps of the video frames needed to be converted to 32-bit integers and be repeated 256 times to construct matching vectors. Thus, a *ScalarBuffer* source operator was added, that produces a sequence of a scalar value defined in its Value property. In order to dynamically change *ScalarBuffer’s* Value property, an *InputMapping* operator was inserted between *ElementIndex* and *ScalarBuffer*. Within the PropertyMappings parameter collection of the *InputMapping* operator, Name parameter was set to ‘Value’ (and is labeled as Value in Figure 1 D) and Selector parameter was set to ‘Index’ (4 repeats) to ensure that the Value parameter was updated with the Index parameter of the *ElementIndex* combinator whenever a frame was acquired. Depth and Size parameters of the *ScalarBuffer* source were set to S32 and (256,1), respectively, to produce a format- and size matching vector to the Open Ephys timestamp packet. Finally, the *WithLatestFrom* and *Concat* combinators ensured that whenever an Open Ephys timestamp packet arrived, the timestamp vector of the recently acquired video frame was assigned to the packet. This approach allowed us to assign each packet of 256 recorded voltage samples to a video frame (Table 1).

**Table 1.** Relationship between LFP samples, video frames and timestamps. **A** LFP sampling rate set at 2 kHz. **B** LFP sampling rate set at 30 kHz. Linear frame interpolation was done offline after data were acquired to assign unassigned frames.

| Timestamp | Time (ms) | LFP Samples | Video Frame |
| --- | --- | --- | --- |
| 1 | 1-128 | 1-256 | 1 |
| 2 | 129-256 | 257-512 | 5 |
| 3 | 257-384 | 513-768 | 8 |
| 4 | 385-512 | 769-1024 | 12 |
| 5 | 513-640 | 1025-1280 | 16 |
| ... |  |  |  |
| 15 | 1793-1920 | 3585-3840 | 15 |
| 16 | 1921-2048 | 3841-4096 | 59 |

| Timestamp | Time (ms) | LFP Samples | Video Frame | Frame Interpolation |
| --- | --- | --- | --- | --- |
| 1 | 0-9 | 1-256 | 1 | 1 |
| 2 | 9-17 | 257-512 | 1 | 1 |
| 3 | 17-26 | 513-768 | 1 | 1 |
| 4 | 26-34 | 769-1024 | 1 | 1 |
| 5 | 34-43 | 1025-1280 | 1 | 2 |
| ... |  |  |  |  |
| 15 | 120-128 | 3585-3840 | 1 | 3 |
| 16 | 128-137 | 3841-4096 | 4 | 4 |

#### Periodic saving of data

*PublishSubject* subject operators were inserted (renamed as ‘Timestamp’, ‘Video’, and ‘Amplifier’) that streamed data to the corresponding *SubscribeSubject* subject operators that were nested into time-dependent loops (Figure 1 D, right). In each loop, *a Timer* source was inserted that produced an output after a user-defined time, that is received by the *TakeUntil* combinator which triggered sink operators to save data. We set DueTime parameter of the *Timer* source to one hour. After the trigger was sent, the *Repeat* combinator reset the *Timer* source, and the process repeated. Video was saved by the *VideoWriter* sink in avi format. Timestamps were saved in 32-bit unsigned integer binary files. Voltage data of eight channels (two from each of four mice) were saved in 16-bit unsigned integer binary files. For naming output files, the suffix property of the *VideoWriter* and *MatrixWriter* sinks was set to the ‘Timestamp’ option and were amended to either ‘vid’, ‘amplifier’, or ‘ts’. Output file naming is important for later GUI analysis.

### Graphical user interface (GUI) for behavioral seizure detection and scoring

Seizure Analysis GUI was developed in MATLAB R2021b and R2022a to analyze seizure activity based on LFP and behavior. The GUI was designed for data organization, seizure detection and seizure scoring purposes.

#### Read and scale voltage data

Voltage data is saved in 16-bit unsigned integer binary files as one long vector. Therefore, in MATLAB data was reshaped based on the number of recorded channels which was determined from the user written config file (see GitHub for details on config file creation - https://github.com/EwellNeuroLab/Seizure-Analysis-GUI). Voltage values are converted to µV by multiplying by a 0.195 µV/bit factor.

#### Seizure detection

LFP data were down sampled (500 Hz) and band-pass filtered (3-50 Hz). The mean activity was then calculated, and peaks were detected that exceeded the user-defined threshold based on mean+N*std, where N is an integer. Next, bursts of spikes (peaks that exceeded the threshold) were identified by grouping consecutive peaks that fired at a minimum frequency of 3 Hz. Burst events were merged within a 2.5 second time window. Finally, epileptic seizures were selected based on the user-defined minimal duration. Events exceeding +/-2000 μV were discarded to reduce the detection of false positives that may occur as a result of mechanical noise.

### Experimental design and statistical analyse*s*

#### Statistical analysis

Statistical analysis was performed in MATLAB 2022a. For comparisons between two groups of data, Pearson-correlation coefficients were calculated between hour-long segments using the *corrcoef()* function and the empirical cumulative distribution was calculated by the *ecdf()* function. For comparisons of three groups, one-way ANOVA was applied using the *anova1()* function. Tukey’s multiple comparison was used for post-hoc analysis with the *multcompare()* function.

#### Software and code

Our Bonsai workflow requires the Bonsai framework that is freely available on https://bonsai-rx.org/. Specific additional packages are required; detailed dependency can be found on our GitHub page. The seizure analysis GUI requires MATLAB 2021b and Signal Processing Toolbox.

#### Code accessibility

The Bonsai workflow for chronic video-LFP recordings and the MATLAB GUI for seizure detection and scoring can be found at https://github.com/EwellNeuroLab.

## RESULTS

Longitudinal recordings were made from mice (n = 6 male, n = 8 female) (Figure 1 B and 1C). Mice were induced with epilepsy using the supra-hippocampal KA model (Bedner et al., 2015) and implanted with two depth electrodes targeting left and right hippocampus, and a ground electrode placed in cerebellum. Immediately after surgery, mice (maximum of four per group) were plugged into one Open Ephys acquisition system for 48 hours to capture the local field potential (LFP) dynamics and video recorded behavior of *status epilepticus* and subsequent disease progression. For the next six weeks, mice were recorded for 48 continuous hours per week.

### Data acquisition and synchronization

Video and LFP data streams were synchronized within Bonsai (Lopes et al., 2015). Sequences of images were acquired from DirectShow multimedia by the *VideoCaptureDevice* source in the Bonsai dataflow (Figure 1 D). We found that the exposure time of our web camera was dependent on the amount of light illumination, which caused variations in frame rates across our 24-hour recording cycle which comprises 12-hour light/dark periods. Therefore, in order to maintain a constant frame rate independent of light condition, auto-exposure of the web camera was disabled within the CaptureProperty parameter of the *VideoCaptureDevice* source. LFP data from the Open Ephys board was acquired by the *RHD2000EvalBoard* source operator (see methods for details).

A crucial step of the workflow was to synchronize the LFP and video data streams (Figure 1 D, left). The Open Ephys acquisition board buffers 256 sampling points of voltage data and sends timestamp in 32-bit integer format. Therefore, the timestamps of the video frames needed to be converted to 32-bit integers and be repeated 256 times to construct matching vectors. Using several bonsai operators we were able to reshape the video data and assign each packet of Open Ephys voltage data to a video frame (see methods for details). In order to quantify the accuracy of our synchronization method, we measured the number of consecutive video frames that were not assigned to the LFP data. We found on average four consecutive frames were unassigned regardless of the LFP sampling rate (Table 1). With a ∼ 30 Hz frame rate, this means that ∼ 130 ms of video data were skipped for every frame that was properly synchronized. Given that for high sampling rates of LFP (> 8000 Hz) each video frame of a 30 Hz camera should be assigned, we concluded that buffering and merging of the video timestamps may be a time limiting factor. However, for purposes like characterizing the behavioral stage of a seizure, this effect is negligible as seizure behaviors happen on the order of seconds and not catching the first 130 ms would not affect the ability to classify Racine score. In case of the need of high time resolution, off-line interpolation of the video timestamps would help mitigate this problem (Table 1B).

In chronic monitoring experiments, data accumulates over multiple days, resulting in large output files that can preclude analysis due to memory and processor limitations. By default, when using Bonsai, data is saved only when the recording is stopped. We therefore extended the workflow to save data regularly during the monitoring (Figure 1 D, right) (see methods for more details). Using subject operators, we were able to implement independent dataflow branches that run parallel with the data acquisition branch; therefore, the frequent data saving does not interrupt video and electrophysiology acquisition. In testing, we found that 637 of 638 hour long files were recorded continuously without any gap between consecutive hour long segments (in one case we found a single missing data packet and lost 256 samples of voltage data), confirming the efficiency of the workflow. Simultaneous monitoring of several animals might require more than one camera. Thus, a Bonsai workflow for correlating two video cameras can be integrated into our workflow, as has been done by others (Lopes & Monteiro, 2021). In that case, whenever a frame arrives from camera A, it is merged with the frame of camera B by the *WithLatestFrom* and *Concat* combinators (https://github.com/EwellNeuroLab/Chronic-Recordings-Bonsai-Workflow).

### Independence of LFP in multi-animal recordings

Simultaneous multi-animal recordings with one Open Ephys acquisition system has not been reported previously to our knowledge, therefore we wanted to verify that there was no bleed-through across channels that would invalidate our data (i.e., if signals from mouse 1 distorted signals from mouse 2). To determine the independence of LFP signals, we calculated correlation coefficients between LFP signals recorded simultaneously between 4 mice (Figure 2 A, two example mice shown). As expected, simultaneous recordings from the left and right hemispheres from the same mouse had moderate levels of correlation due to periodic bouts of synchronous activity across the left and right hemispheres (Figure 2 B & C, grey) (0.24 ± 0.02, n = 96 comparisons). In comparison, signals recorded simultaneously from different mice (Figure 2 B & C, purple) had negligible near zero correlation (1.35*10^−5^ ± 0.16*10^−5^, n = 288 comparisons), which was similar in magnitude to chance correlation (Figure 2 B & C, cyan), calculated from comparisons between non-simultaneous recordings from different mice (7.75*10^−6^ ± 0.65*10^−6^, n = 288 comparisons). There was a significant group effect (One-Way ANOVA, F(2,672) = 344.7, *p* = 9.5*10^−104^), with post-hoc comparisons revealing a significant difference between the left-right comparison and both the simultaneous recordings across animals (*p*<0.001) and the comparisons across animals and hours (Figure 2 C, cyan, *** *p*<0.001) but not between the latter two. These results suggest that the signals from individual mice do not interfere with each other.

**Figure 2.**
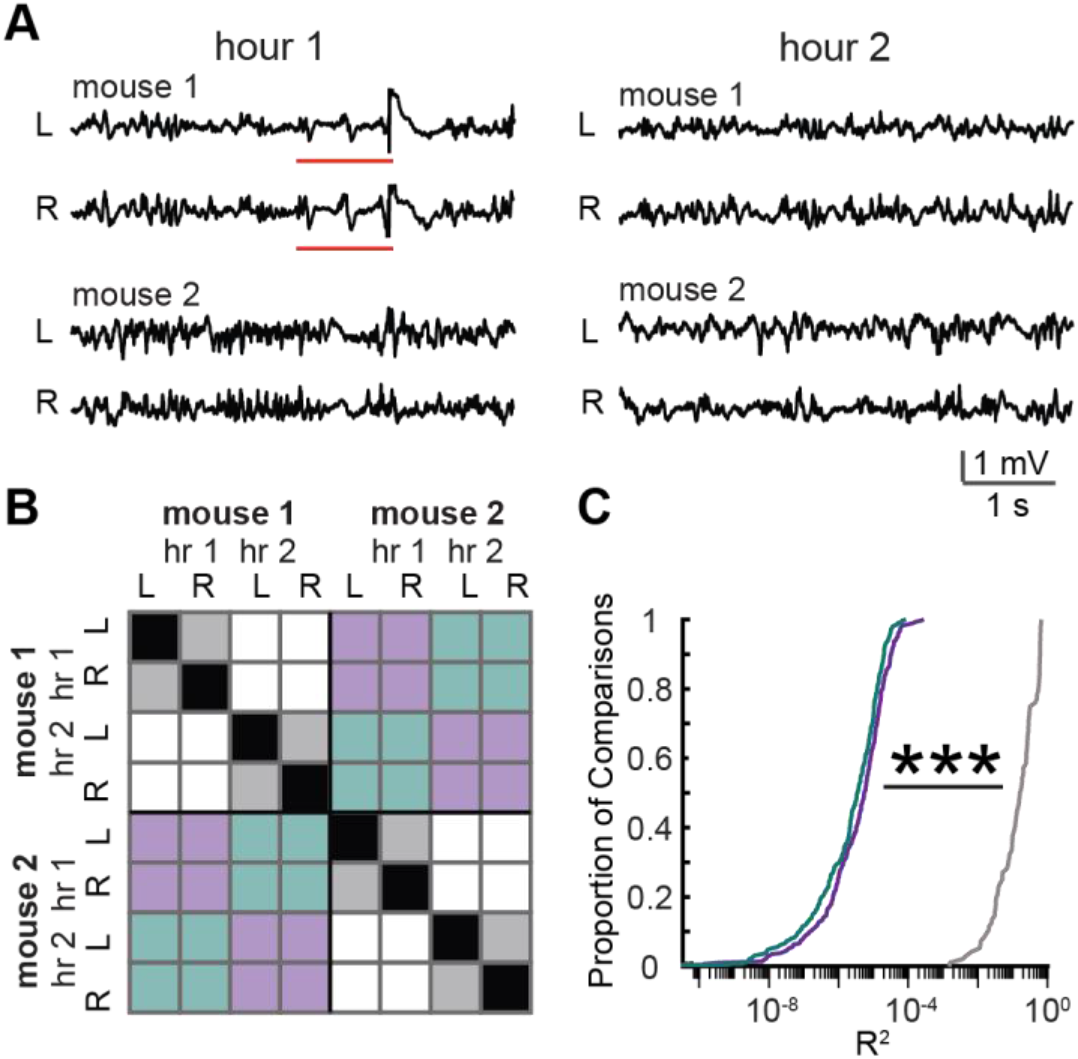
LFP signals simultaneously recorded from different mice are independent. **A** Five-second-long clips of LFP recorded simultaneously from two mice during hour 1 of monitoring (left) or hour 2 of monitoring (right). For each mouse two traces are shown corresponding to electrodes located in left (L) and right (R) hippocampus. Red lines indicate a period in which signals from both hemispheres showed synchronous signal within mouse 1. **B** A matrix to visualize comparisons made to create distributions shown in (C). Correlations between hour long traces were made for comparisons that fell into three categories: (1) left vs right same mouse, same hour – grey; (2) different mice, same hour (purple); (3) different mice, different hours (cyan). Black represents self-comparisons, which were not included. White represents comparisons between the same mouse, but different hours, which were also not included. For actual distributions shown in (C), comparisons were made across 4 mice, and for each hour across 24 hours of recording. **C** Cumulative Density Functions (CDFs) of correlations obtained from comparisons described in (B). *** *p* < 0.001, One-Way ANOVA, post-hoc Tukey.

### Seizure scoring with the Seizure Analysis GUI

Identifying epileptic seizure LFP events and correlating them with behavior is time consuming. Thus, a MATLAB graphical user interface (GUI) was developed to provide a fast and reliable way to analyze seizures without any programming experience required by the user. The GUI allows the user to handle large datasets, visualize LFP and video, and detect and score seizures (Figure 3 A).

**Figure 3.**
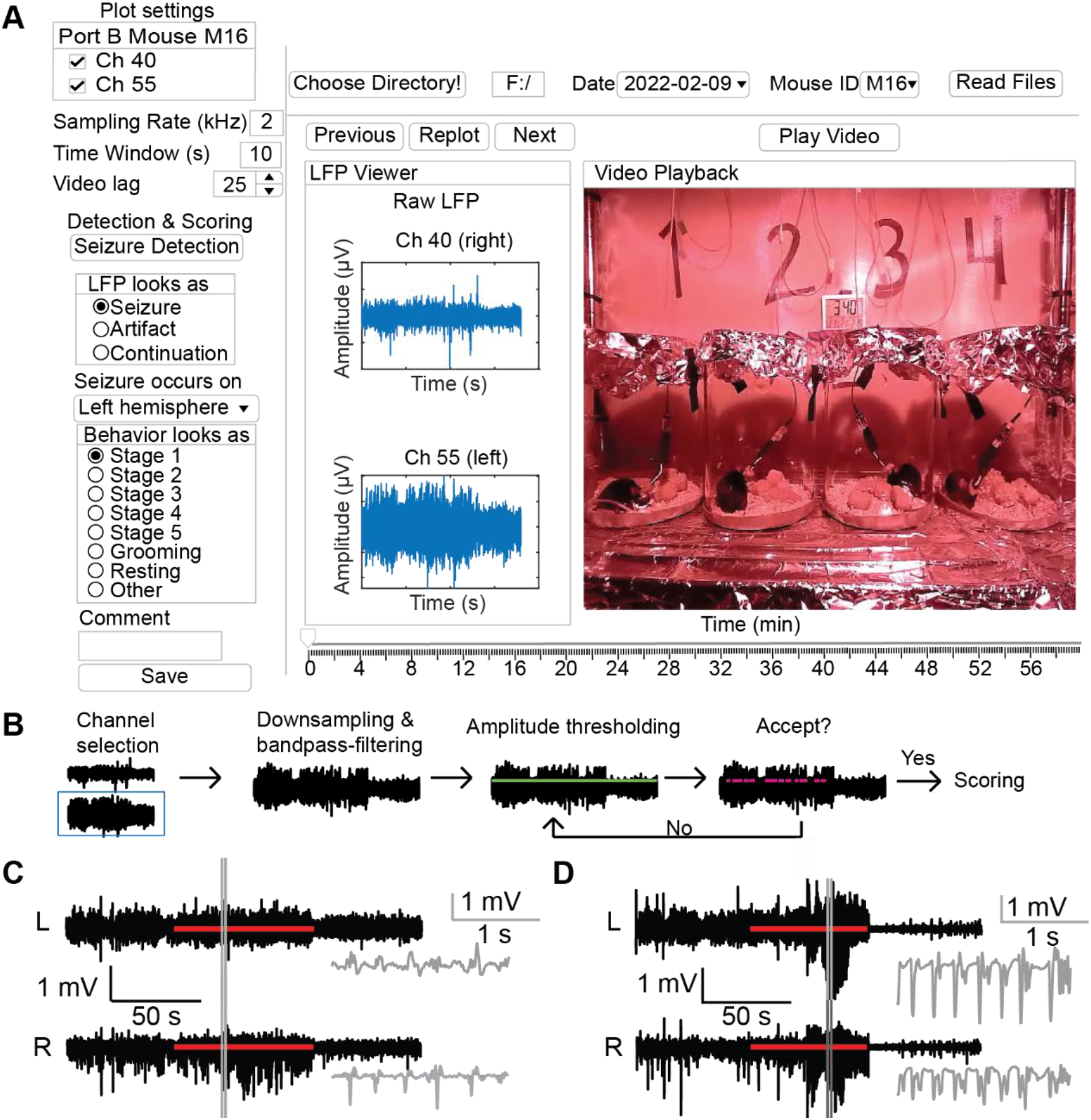
Seizure Analysis GUI. **A** Schematic of the GUI. A working directory is selected and recording date and mouse are chosen to be analyzed (top, right). Channels to show and duration of LFP traces are set by the user in Plot settings (top, left). Video lag, allows the user to set playback speed. Pressing ‘Seizure Detection’ opens a pop up window where the user may filter and/or flip signals before applying a threshold (B). Putative seizures will be highlighted in magenta, allowing the user to decide to proceed or redetect with different parameters. When the user accepts the detection results, video and filtered LFP of the first event is loaded into the Video Playback and LFP Viewer. Detected seizure events are toggled through by the user using the ‘Next’ button. Questions regarding the LFP signal and behavior are answered for each event (left). After performing scoring, annotations are saved by pressing the ‘Save’ button. **B** Seizure detection sequence. The user selects a channel on which the detection will occur (blue box). LFP is down-sampled, bandpass-filtered, and possibly flipped according to user input. An amplitude threshold is set by inputting an integer value (green line). Finally, detected events are displayed (magenta). User can expand view to look at several individual events and then choose to accept all results and start scoring or adjust the threshold and repeat the detection. **C** Example of detected subclinical seizure. Detected seizure is indicated by the red bar and additional 60 s pre- and post-seizure baseline are shown. Grey lines indicate the range of the zoomed in 2 s long gray trace. In this example, the seizure is only observed on the right hemisphere (bottom). **D** Example of stage 5 seizure. Seizure can be observed on both hemispheres. Notations are the same as in (C).

#### Data organization

Using the Bonsai workflow for chronic recordings, each hour of the recording saved a video file, timestamp file, and voltage data file. The first step is to select a working directory where the Bonsai output files can be found. We structured our data such that consecutive hours over the 48-hour recording period were all written to the same directory. Voltage files contain recorded data from the four mice. An additional configuration file must be provided in the working directory that contains information about the mice and recorded channels, to aid in parsing the voltage file. Based on the information from the configuration file and date and time of acquisition, the user selects and loads data of a specific recording and mouse for further analysis.

#### Visualization

Once the data is loaded, LFP and corresponding video are visualized in two separate windows. The user may select specific channels to view and set a time window for LFP and video replay, respectively. Video replay speed may vary among computers based on performance; therefore, a *Video Lag* variable and a calibration method is provided to ensure the real-time replay of the video.

#### Seizure detection

Seizures will be detected on a user selected LFP data channel (Figure 3 B). The user can select boundaries of a bandpass filter, a down sampling rate, a signed multiplier (for flipping the signal), and a detection threshold. These settings are saved and can be loaded back into the GUI to recall previously used settings. Putative seizures are detected based on the algorithm described in the Materials and Methods section. The user may decide to accept the detected events or change the setting parameters. Note that the algorithm is suitable for detecting epileptic seizure events for scoring and behavioral correlation, but not applicable for analyzing LFP dynamics at the on- and offset of seizures because the algorithm has not been optimized for determining exact on- and offsets. If this is the aim of the user, we recommend using the timestamps outputted by our GUI to flag seizure events, and reanalyze those LFP periods using additional analytical tools.

#### Seizure scoring

The user may toggle between detected events to visualize and score events one by one. The corresponding video around each putative seizure event is played in pre-seizure - seizure – post-seizure segments in the Video Viewer. Pre- and post-seizure duration can be set in the Plot settings module. When the video is playing, a moving color-coded bar on the LFP Viewer represents the elapsed time. After viewing, the user answers a series of questions to classify the event. Question one asks to classify the LFP as a true or false positive seizure event. False positive events may be detected due to mechanical noise associated with repetitive movements of the mouse, such as grooming or eating. Once classified as a true positive, the user will have the option to mark the seizure as a continuation of a previous event or as a new event. Question two asks the user whether the seizure is observed on the left, the right, or both hemispheres. Finally, question three asks for the seizure score based on the Racine scale (Stage 1-5) (Racine, 1972). Any additional observations can be described in a comment field. Event characteristics (timing of onset and offset, duration of seizures) and scoring parameters (LFP and behavior classification, and any additional comments) are saved in a summary MATLAB file for further analysis and statistical purposes. Each output file is named with a user identifier and date of scoring, allowing the analysis of the same file by several people, which is ideal for facilitating blind scoring by several users.

Our GUI was successfully applied for detecting and scoring epileptic events by several undergraduate researchers. We were able to observe and score subclinical and behavioral seizures (Figure 3 C & D). Although our GUI is customized for epileptic seizure analysis, replacing the seizure detection function allows one to repurpose the software for distinct applications.

## DISCUSSION

When studying epilepsy, simultaneous monitoring of the neurophysiological signals and behavior is necessary to make meaningful connections between the observed neural activity and the subject’s behavior (Berger et al., 1929). Many groups have developed various software packages and platforms for managing neurophysiological and video data (Campagnola, Kratz, & Manis, 2014; Black et al., 2017; Kimchi et al., 2020). Here, we aimed to develop a platform that would allow for the flexibility to perform simultaneous LFP and video monitoring, as well as high density single unit recordings which are becoming more common in epilepsy research (Lenck-Santini & Holmes, 2008; Hernan et al., 2014; Ewell et al., 2015; Ewell et al., 2019; Dahal et al., 2019; Shuman et al., 2020). For multi-animal LFP-video recordings, we constructed home-made depth electrodes and wired them to an Intan adapter board to facilitate LFP recordings (2 channels per animal) in 4 animals simultaneously. Wiring to an Intan adapter board makes our system compatible with virtually any home-built recording device (<16 channels) that could similarly be wired to an adapter board such as custom-designed 3D printed headplates shown previously to enable reliable recordings in epileptic mice (Zhu, Aiani, & Pedersen, 2020). Simply switching headstages allows for recordings of up to 128 channels per animal and is compatible with the rest of our workflow.

When choosing which software to implement our workflow, Bonsai was the natural choice, as there was already an open-source Bonsai module available for streaming LFP data acquired with an Open Ephys acquisition system. We wanted to implement a basic workflow that could offer maximal flexibility to future experimental designs. For example, we could imagine real-time behavioral scoring of seizures by adding behavioral tracking and pose estimation analysis to the video stream, as has been already implemented in Bonsai in other workflows (Dreosti et al., 2015). Furthermore, our workflow could easily be updated to realize chronic monitoring of other data types such as fiber photometry (Soares, Atallah & Paton, 2016), calcium imaging (Sotskov et al., 2022), and high-density electrophysiology (Karalis & Sirota, 2022). Finally, Bonsai has been used in different animal models, including zebrafish and drosophila, suggesting that our platform may be translatable to other animal models of epilepsy (Itskov et al., 2014; Hortopan, Dinday, & Baraban, 2010; Sun et al., 2012).

Bonsai is ideal for controlling the timing/triggering of external hardware (i.e., laser pulses for optogenetics) and is compatible with microcontrollers, such as Arduino, providing a vast toolbox for the experimenter (Douglass et al., 2017). We chose a data stream synchronization method that can be performed with an inexpensive web camera but has limited temporal resolution, due to the buffering of the video timestamps and the low frame rate of the camera. When high precision timing is needed, as in many closed-loop experiments, the best solution would be to invest in a higher frame rate camera that sends shutter events to Bonsai (Buccino et al., 2018). However, due to the reactive nature of Bonsai, when a global event occurs (i.e., triggering a laser), timestamps of the recently acquired camera frame and LFP packet can be saved individually and thus without buffering, creating a single, synchronized video-LFP timestamp for that specific event. Therefore, precise records can be made, even with our low-cost solution.

We developed the Seizure Analysis GUI that incorporates a seizure detection algorithm based on frequency and amplitude properties of the occurring spikes in the LFP. The algorithm may detect false positive events, which must be corrected by the user during seizure scoring. Our analysis did not require the accurate timing of on- and offset of the seizure and therefore the algorithm has not been optimized for this purpose. Several seizure detection algorithms have been developed that can be used for precise analysis of seizure dynamics (Krook-Magnuson, Armstrong, Oijala, & Soltesz, 2013; Richard et al., 2015; Pfammatter et al., 2018; Andrade et al., 2018). When needed, the source code of the GUI can be modified and the seizure detection algorithm can be replaced with an alternative algorithm that is ideal for the user, while all the other features of the GUI remain.

The recent explosion of open-source hardware, software, and data in neuroscience is pushing the limits of neuroscience by making it more accessible and easier to share (Gleeson et al., 2017). We have developed our multi-animal chronic monitoring platform in this spirit, and have worked to make it as low-cost as possible, all while maximizing the flexibility that will allow other laboratories to invest in one acquisition system that would support many types of neurophysiological experiments.

## Acknowledgements

We would like to thank the American Epilepsy Society for funding L.A.E. and UCI EpiCenter for NINDS T32 to B.L.B. We would like to thank Cathy B. Dang, Usean J. Redic, and Ramy Labib for testing our seizure detection and analysis GUI and providing feedback.

